# Heterogeneous pro-inflammatory response to BRAFV600E-induced thyroid tumor development

**DOI:** 10.64898/2026.03.26.714444

**Authors:** Sima Kumari, Carmen Moccia, Henrik Fagman, Elin Schoultz, Mikael Nilsson

## Abstract

**Background:** The tumor immune microenvironment likely plays a central role in progression of thyroid cancer. As for most other solid tumors, it is unknown if immune dysregulation contributes to earlier, subclinical stages of thyroid tumor development, or whether thyroid tumor heterogeneity might involve differential expression of pro-inflammatory mediators.

**Methods:** The time course of tumor-associated inflammation was studied in *Tg-CreER^T2^;Braf ^CA/+^* mice representing a model of BRAF^V600E^-driven papillary thyroid carcinoma (PTC). Tumor growth was estimated by histological examination and magnetic resonance imaging. Cytokine expression was monitored by quantitative RT-PCR, RNAScope and Western blot analyses.

**Results:** Based on spontaneous *Braf^CA^* activation due to leaky Cre activity in a minority of targeted cells tumors developed within a preserved thyroid tissue architecture to multifocal papillary thyroid carcinoma (PTC) over a period of 12 months. Tumorigenesis was accompanied by a gradually increased mRNA and protein expression of interleukin-1beta (IL-1β), interleukin-6 and tumor necrosis factor-alpha (TNF-α) starting already before Braf mutant cells commenced neoplastic growth. RNAScope revealed that both follicular cells and stromal cells expressed *Il1b* whereas *Il6* and *Tnfa* transcripts were mostly confined to neoplastic epithelia. Early cytokine expression was associated with oncogene-induced senescence, whereas during tumor development (3-6 months) and in advanced tumor stages (at 12 months) the cytokine expression pattern differed among glands and tumor foci of the same gland accompanied by a highly variable locoregional lymphocytic infiltration. Oral treatment of mutant mice for 1 month with PLX4720, a vemurafenib prodrug, partially reduced cytokine expression along with inhibited tumor growth and redifferentiation of thyroid function. The magnitude of reduced cytokine expression differed much between glands and among mice of both sexes.

**Conclusions:** These findings indicate that oncogenic BRAF^V600E^ targeted to the thyroid both stimulates endogenous production of IL-1β, IL-6 and TNF-α and recruits inflammatory cells to foci of early tumor development. PTCs of different clonal origin are distinguished by differential expression of pro-inflammatory cytokines. The anti-inflammatory effect of mutant Braf kinase inhibition varies presumably related to heterogeneous tumor development, which evolves from stochastic *Braf^CA^* activation suggesting there are clonally different probabilities of acquiring drug resistance among Braf mutant thyroid follicular cells.

## Background

Cancer-associated inflammation is a prominent feature of thyroid carcinoma (1–3). Most studies so far have been characterizing the presence of infiltrating immune cells and expression of cytokines and chemokines in biopsies of manifest tumors or cell lines established from various thyroid cancer types (4, 5), inferring a likely pathogenic role of pro-inflammatory mediators in tumor progression (6). However, immune therapy is yet of limited positive value as a potential treatment option for patients with advanced thyroid cancer (7). Putative influences of the tumor immune microenvironment in earlier stages of thyroid carcinogenesis are unknown due to suitable *in vivo* models of thyroid tumor initiation and tumor development are lacking.

Oncogenic drivers may be conditionally expressed in the mouse thyroid to reproduce development of the corresponding tumors in humans. Hence, inducible thyroid targeting of a mutant Braf allele encoding BRAF^V600E^, the most prevalent driver mutation in thyroid cancer, gives rise to tumors that strongly resemble papillary thyroid carcinoma (PTC) in humans (8). However, a major confounding factor of this model is that Braf oncoprotein is synchronously expressed in most if not all thyroid cells with little if any normal thyroid tissue remaining (9). Moreover, since *Braf^CA^* activation constitutively activates the MAPK signaling pathway the affected thyroid cells are functionally dedifferentiated leading to severe hypothyroidism (10), which in turn triggers goitrogenic growth that overrides tumorigenesis and makes it difficult to distinguish neoplastic from reactive responses. Notably, recent findings infer an immunomodulatory role of the thyroid follicular epithelium by contact-dependent inhibition of T-cell proliferation (11, 12). Although an immunosuppressive effect primarily might be of importance for the risk of developing autoimmune thyroid disease, this opens for a possibility that immunosurveillance of oncogenic mutant cells might be influenced by their normal follicular cell counterparts. Thus, from both locoregional and systemic viewpoints currently available models of induced oncogene activation appears unfavorable when it comes to exploration of immunological factors and immunomodulatory effects that might be implicated in sporadic tumor development in the thyroid gland.

We and others have discovered that thyroid carcinomas spontaneously develop in *TgCreER^T2^*;*Braf^CA/+^* mice mediated by leaky activation of Cre recombinase (13, 14), which in the absence of the inducing agent (tamoxifen) infers stochastic activation of a *Braf* mutant allele encoding BRAF^V600E^ oncoprotein in a limited number of thyroid follicular cells (15). In this model, multifocal tumors develop in euthyroid conditions and allows monitoring of tumor clonal origin and heterogeneous growth patterns evolving into histologically distinct PTC subtypes (14, 16). Moreover, tumor development is oligoclonal and influenced by intrinsic properties of the originating follicle that appears to be an independent determinant of clonal selection promoting or antagonizing tumor growth (14). Here, employing the sporadic mouse PTC model we pinpoint spatiotemporal expression changes of major pro-inflammatory cytokines during BRAF^V600E^-driven thyroid tumorigenesis – from tumor initiation to advanced tumor stages –and investigate the anti-inflammatory response to mutant Braf kinase inhibition.

## Methods

### Mouse strains and experiments

*TgCreER^T2^* mice (RRID:IMSR_JAX:030676; The Jackson Laboratory) with inducible Cre under control of the *Tg* promoter (17) were crossed with *Braf^CA^* mice (RRID:IMSR_JAX:017837; Jackson) heterozygous for mutant *Braf* encoding BRAF^V600E^ (18) to generate *TgCreER^T2^;Braf^CA/+^*mice, as previously reported (14). Strains were backcrossed with C57BL/6J female mice (B6-F; Taconic Biosciences) 10 times before recombination. Ear punch biopsies were sampled for genotyping with PCR. Tamoxifen dissolved in sunflower oil (10 mg/ml) was injected intraperitoneally (50 µl) to pregnant mice twice, 14 and 15 days post-coitum, for CreER^T2^ induction accompanying embryonic onset of thyroglobulin expression. Age at endpoint of experiments were 10 days (d), 30 d and 3-6-7-12 months (mo). Oral PLX4720 (417 ppm; provided by Plexxikon), a prodrug to vemurafenib, and control dietary pellets were continuously supplied during treatment period. After sacrifice and photo-documentation of *in situ* gland size, thyroids were excised *en bloc* with the trachea and esophagus, fixed in 4% paraformaldehyde and embedded in paraffin after which sections were subjected to routine hematoxylin–eosin staining and immunostaining (see below), or sampled free from extrathyroidal tissue for mRNA and protein analyses. Animal housing and experiments were approved by Göteborgs Djurförsöksetiska Nämnd (5.8.18-04502/2023) according to European standards and national regulations provided by the Swedish Board of Agriculture. The ethics permit is valid until May 17^th^, 2028, comprising 8000 individuals; in total 164 mice were used in the current animal study, which started in May 2023 and ended in August 2025. Animals were kept at the animal facility of Sahlgrenska Academy, the Laboratory of Experimental Medicine, with standard pellet diet and sterile water ad libitum, unless otherwise stated (drug treatment and live imaging procedure). Phenotypes of mutants with or without drug treatment were compared with wildtype mice of the same or parallelly bred litters. Each experimental setting (animal age, duration of experiment, induced vs non-induced, drug-treated vs untreated and type of analysis) were investigated at least twice and mostly more than three times. Sample size (n/group) was in a sense depending on breeding outcome. Male and female mice were included in each experimental group of comparison i.e. inclusion of animals was not randomized. Analysts were primarily blinded to genotypes of investigated tissue samples except for western blot analysis. Exclusion criteria were solely on basis of sudden illness or death that very rarely occurred even in tumor-bearing mice of high age (12 months or older). Humane endpoints comprised weight loss, deteriorated condition or respiratory distress. However, no adverse health effects related to experimental tumor growth were noticed, hence no data points were excluded from analysis. Method of euthanasia comprised asphyxiation with carbon dioxide in accordance with veterinary guidelines. Animal numbers subjected to each analysis are given in figure legends.

### Quantitative real-time PCR (qPCR) analysis

Thyroid samples stored at −80°C in RNA*later*™ solution (Thermo Fisher Scientific) were homogenized using a TissueLyser II (Qiagen) and RNA-extracted with RNeasy mini kit (Qiagen). RNA amounts and quality were determined spectrophotometrically (NanoDrop1000; Thermo Scientific). cDNA was synthesized using TATAA GrandScript cDNA Synthesis Kit (TATAA Biocenter) and T100 Thermal Cycler (Bio-Rad) and stored at −20°C. Primers to mouse *Tg* (forward: 5′-CATGGAATCTAATGCCAAGAACTG-3′; reverse: 5′-TCCCTGTGAGCTTTTGGAATG-3′), *Tpo* (forward: 5′-CAAAGGCTGGAACCCTAATTTCT-3′; reverse: 5′-AACTTGAATGAGGTGCCTTGTCA-3′), *Tshr* (forward: 5′-TCCCTGAAAACGCATTCCA-3′; reverse: 5′-GCATCCAGCTTTGTTCCATTG-3′), Slc5a5 (forward: 5′-TCCACAGGAATCATCTGCACC-3′; reverse: 5′-CCACGGCCTTCATACCACC-3′), *Pax8* (forward: 5′-GATAGGAGACTACAAGCGGCA-3′); reverse:5′CGGATGATTCTGTTGATGGAGC-3′), *Il1b* (forward: 5′-CCTTCCAGGATGAGGACATGA-3′; reverse: 5′-TGAGTCACAGAGGATGGGCTC- 3′), *Il6* (forward: 5′-GATACCACTCCCAACAGACCT-3′; reverse: 5′-CTCATTTCCACGATTTCCCAGA-3′), *Tnfa* (forward: 5′- TGCCTATGTCTCAGCCTCT-3′; reverse: 5′-GAGGCCATTTGGGAACTTCT-3′) - were designed in Primer-BLAST (19) based on sequences from public databases (Ensembl or Santa Cruz Genome Browser). BLAST analysis verified that the selected primer sequences were sufficiently different from the rest of the mouse transcriptome. *In silico* oligonucleotide secondary structure prediction was performed with NetPrimer (PREMIER Biosoft International). Specificity of primers was confirmed by agarose gel electrophoresis of amplicons. A reference gene panel for mouse (TATAA Biocenter) was evaluated with NormFinder; *Gapdh* was chosen as optimal reference gene on basis of predefined software criteria for relative quantification. Quantitative real-time PCR (qPCR) was performed using a CFX384 Touch real-time cycler (Bio-Rad) and TATAA SYBR GrandMaster Mix (TATAA Biocenter), 400 nM primer and 2 μl cDNA. Expected PCR products were confirmed by agarose gel electrophoresis and all samples were analyzed by melting-curve analysis.

### In situ hybridization (ISH) by RNAscope

ISH was conducted using the RNAscope® 2.5 HD Detection Reagents-RED (Advanced Cell Diagnostics, 322360). Briefly, formalin-fixed paraffin-embedded (FFPE) sections (5 μm thick) were cut and collected on SuperFrost Plus® Slides (Epredia; Cat. No. J1800AMNZ) and dried at 60°C for 1 hour. Tissue sections were subsequently deparaffinized with xylene (2 × 5 minutes) and absolute ethanol (2 × 1 minute) at room temperature, followed by a 10-minute incubation with hydrogen peroxide. Epitope was retrieved by exposing the slides to RNAscope 1× Target Retrieval Reagent (Advanced Cell Diagnostics) for 30 minutes at 98°C. Tissues were then permeabilized using RNAscope protease plus (Advanced Cell Diagnostics) for 30 minutes at 40°C, followed by the RNAscope™ Probe (*Mm-Tnfa*, cat No. 311081, *Mm-Il1b*, cat No. 316891, *Mm-Il6*, cat No. 315891) hybridization was carried out at 40 °C for two hours, followed by sequential hybridizations with the preamplifier, amplifier, and label probes to target RNA molecules. RNAscope probes for the housekeeping gene *Ppib* and bacterial gene *dapB* were used as positive and negative controls, respectively. Digital whole slides were obtained using a wide-field Zeiss Axioskop2 plus microscope equipped with a Nikon DS-Qi1Mc camera was used for imaging.

### Western blot (WB) analysis

Mouse thyroid tissues were processed using tissue protein extraction according to the manufacturer’s protocol (Thermo Fisher Scientific). After centrifugation at 16,000□×□g for 5□min, the protein in the supernatant (cytoplasmic extract) was transferred to a prechilled tube. Insoluble fraction (pellet) was suspended in an ice-cold nuclear extraction reagent, vortexed, and centrifuged at 16,000□×□g for 10□min. WB analysis was performed after measuring the protein concentrations using the BCA Protein Assay Kit (Thermo Fisher) to ensure equal loading of the protein. After the transfer of the proteins to the membrane (Trans-Blot Turbo Midi Nitrocellulose Transfer Membrane; Bio-Rad), the membranes were colored using Ponceau (Thermo Fisher) to verify that the transfer procedure was successful. Specific cytokines proteins were detected using primary antibodies against tumor necrosis factor-alpha (anti-TNF-α; Abcam, Cat no. ab6671), interleukin-6 (anti-IL-6; Abcam, Cat no. ab208113) interleukin-1 beta (anti-IL-1ß; Abcam, Cat no. ab9722), P21 (anti-P21; Abcam, Cat. no. ab188224) and β-Actin (anti-beta-actin/ACTB; Sigma-Aldrich, Cat. no. A5441). Specific bands were detected by enhanced chemiluminescence following labeling with HRP-linked anti-rabbit (Cell Signaling, Cat no. 7074S) and anti-mouse (Cell Signaling, Cat no. 7076S) secondary antibodies and use of PageRuler^TM^ Plus Prestained Protein Ladder (Thermo Fisher, Cat. no. 26619). Protein levels were analyzed by densitometry. Representative WBs are shown with β-actin used as loading control. Data are mean ± SEM from n=3 per group; *P < 0.05, **P < 0.01, ***P < 0.001).

### Immunohistochemistry (IHC)

Deparaffinized 4 um thick sections were subjected to epitope retrieval by PT Link (Dacocytomation) and quenching of endogenous peroxidase activity prior to immunostaining optimized with the Dako EnVision system. Antibodies against P21, IL-1ß, IL-6 and TNF-α were identical to those used for western blotting. Sections were viewed and imaged as detailed above for ISH. Notably, IHC of the indicated cytokines did not result in any reliable immunostaining and was therefore omitted from presentation.

### Magnetic resonance imaging

Anatomical magnetic resonance (MR) T2 weighted neck scans were acquired using a 7T horizontal bore pre-clinical MR system with a 72-mm volume coil (Bruker BioSpin MRI GmbH, Germany; software: ParaVision 5.1), and a 4-channel array rat brain receiver coil for signal reception (RAPID Biomedical GmbH, Germany). Live animals were imaged in prone position and kept under isoflurane anesthesia (2-3%, Isoba vet. Schering-Plough Animal-health, Denmark) and surveillance of body temperature and breathing. Individual mice were repeatedly examined with MR at 3, 6 and 12 months of age, which likely excluded monitoring of any hypothetical confounding effect on tumor growth rate post-MR examination.

### Statistical analysis

Statistical significance between the groups was evaluated by one- way ANOVA with t test for post hoc analysis. Differences were considered statistically significant at P < .05. The statistical analyses were performed using SPSS Statistics (v2; IBM Corp., Armonk, NY, USA).

## Results

### Prenatal Braf^CA^ activation triggers excessive interleukin-6 expression in the juvenile mouse thyroid

The initial pro-inflammatory response to global versus stochastic oncogene activation in the thyroid was investigated in *TgCreER^T2^;Braf^CA/+^*mice that were injected or not with tamoxifen. The rationale for comparing these two conditions is that in the absence of tamoxifen induction there is a considerable amount of spontaneous *Braf^CA^* activation in a minority of thyroid cells due to leaky Cre activity which drives multifocal, clonally independent tumorigenesis within a preserved thyroid follicular tissue (Fig. 1A) (14). By contrast, tamoxifen is expected to synchronously activate *Braf^CA^* in most if not all cells that express the Cre driver (thyroglobulin) by which the entire gland will become tumorous (Fig. 1A) (13). Since non-induced activation starts in infancy, possibly already before birth, and much more efficiently generates tumors than if mutant BRAF is expressed in adult and largely quiescent thyroid cells (14, 20), we adopted a protocol in which pregnant *TgCreER^T2^;Braf^CA/+^*mice were injected or not with tamoxifen timing the onset of thyroglobulin expression along with embryonic thyroid differentiation (21), after which the expression of major pro-inflammatory cytokines was monitored by qPCR in thyroid samples obtained postnatally (Fig. 1B, upper panel). As shown in Fig. 1B, induced *Braf^CA^*activation *in utero* markedly stimulated gene expression of *Il-6* and to a lesser extent that of *Il1b*) and *Tnfa*, as evident at P10 and sustaining at P30. By contrast, following spontaneous activation of mutant Braf *Il1b* and *Il6* transcripts were only moderately and equally increased at P30 but not earlier, and *Tnfa* was only slightly increased above the control level (Fig. 1B).

**Fig. 1.**
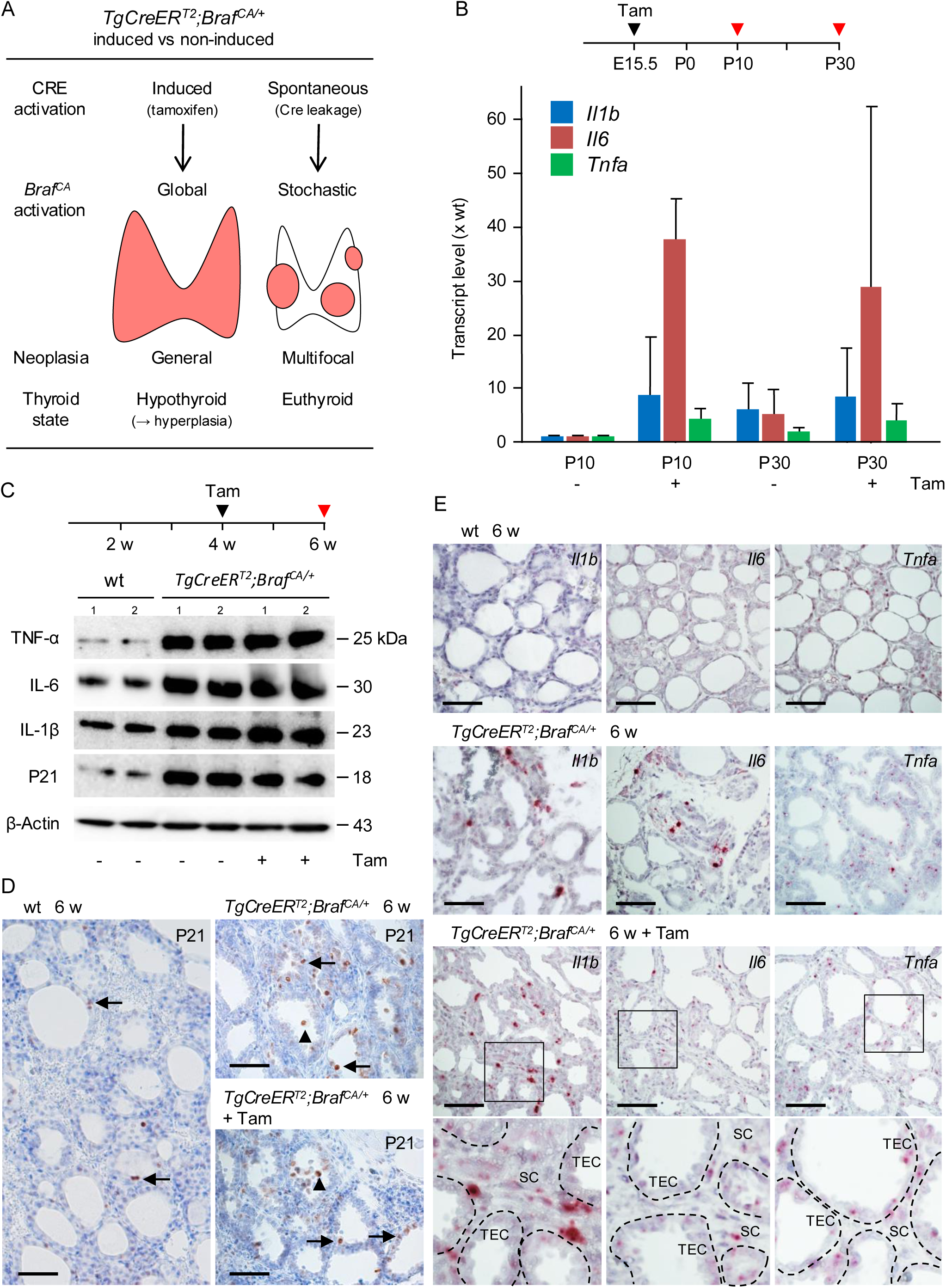
Expression of pro-inflammatory cytokines during BRAF^V600E^-driven early tumor development in mouse thyroid. Thyroid targeting of a mutant *Braf* allele was obtained by CRE recombinase conditionally expressed by the thyroglobulin (*Tg*) promoter in *TgCreER^T2^;Braf^CA/+^* mice. **A** Basic features of the experimental model following induction with tamoxifen or in non-induced conditions. Collectively from: (14–16). **B** Cytokine expression after induced *Braf^CA^* activation in embryonic thyroid. Quantitative real-time PCR (qPCR) analysis. Tamoxifen was injected twice to pregnant mice corresponding to E14.5 and E15.5 i.e. concomitant with onset of *Tg* expression. Mean ± SD (P10, induced: n=2; P10, non-induced: n=3; P30, induced: n=3; P30, non-induced: n=3). Transcript expression levels were compared to that of control thyroids obtained from age-matched non-mutant (wt) mice (n=2 for each time point). **C-E** Induced and non-induced expression of IL-1β, IL-6, TNF-α and P21 in thyroid of young adults. Data collected from samples of the same experiment comprising western blot (WB) (C), immunohistochemistry (D) and RNAScope (E) analyses. Arrows and arrowheads in (D) indicate P21^+^ cells present in the follicular epithelium and inside the lumen, respectively. In (E), lower panel shows insets at high magnification with follicles encircled. wt wildtype, E embryonic day, P postpartum, Tam tamoxifen, IL-1β interleukin-1beta, IL-6 interleukin-6, TNF-α tumor necrosis factor-alpha, TEC thyroid epithelial cells, SC stromal cells. WB: n=3 for all groups; * p<0.05, ** p<0.01, *** p<0.001. Scale bars: 100 µm.

### Spontaneous Braf^CA^ activation upregulates pro-inflammatory cytokines and P21 to the same magnitude as conditionally induced activation in adult mouse thyroid

Next, we investigated whether adult thyroid cells might respond similarly to induced *Braf^CA^* activation in embryonic thyroid development with strongly enhanced IL-6 expression plausibly associated with oncogene-induced senescence (22). For comparison, both induced and non-induced mutant mice were examined 2 weeks after tamoxifen injections (Fig. 1C, upper panel). Remarkably, this showed a blunted inflammatory response that in fact did not differ from the expression levels of IL-1β, IL-6 and TNF-α in non-induced *TgCreER^T2^;Braf^CA/+^*mice (Fig. 1C). Moreover, there were no major difference in number and distribution of P21^+^ cells in induced versus non-induced conditions (Fig. 1D); although more cells became senescent and some follicles showed shedding of P21^+^ cells into the lumen most follicular cells remained P21 negative despite induced oncogene activation.

From this we conclude that effects of spontaneous *Braf^CA^*activation predominates in young adult thyroid cells and that induction does not add much to the mutant phenotype. We previously showed in clonal tracing experiments using a double fluorescent reporter that superimposed induction affects only a minor fraction of cells (14, 16). Two mechanisms probably explain why adult thyroid cells respond poorly to induced activation: an increasing number of cells express mutant Braf due to leaky Cre already before induction and some non-mutant follicular cells may also be dedifferentiated due to a bystander effect, both conferring downregulation of the Cre driver and hence insensitivity to tamoxifen.

### Mutant Braf elicits cytokine expression concomitantly in neoplastic thyroid follicular cells and stromal cells

Since IHC staining of IL-1β, IL-6 and TNF-α with the same antibodies used for Western blotting did not produce consistent and reliable results (data not shown), we decided to identify cytokine-expressing cells by RNAscope ISH assay on FFPE tissue sections (Fig. 1E). In wildtype mice, thyroid tissue showed weak labeling most evidently for *Tnfa* and largely confined to follicular epithelial cells (Fig.1E, upper panel). The number of *Il1b*, *Il6* and *Tnfa* positive thyroid cells increased in 6 weeks old *TgCreER^T2^;Braf^CA/+^*mice in both non-induced (Fig. 1E, middle panel) and induced (Fig. 1E, lower panels) animals. *Il16*^+^ cells mostly accumulated in the stromal compartment whereas *Il6* and *Tnfa* expression comprised both follicular and stromal cells. Cytokine-expressing stromal cells were more frequent after induction than without. Notably, the distribution of cytokine-expressing epithelial cells varied both among and within follicles consistent with follicular heterogeneity and putatively different sensitivity to oncogene activation. Clustering of *Il1b*^+^ cells and to some extent *Il6*^+^ cells around some follicles further suggested that the inflammatory response to BRAF^V600E^ involve local recruitment of cytokine-producing non-mutant cells to the pre-tumorous tissue microenvironment.

### Progressively increased cytokine expression accompanies tumor development to papillary thyroid carcinoma

We and others have previously shown that PTC tumors appear in 12 months old non-induced *TgCreER^T2^;Braf^CA/+^* mice due to stochastic *Braf^CA^* activation which, in contrast to after induction by tamoxifen, occurs in a preserved follicular tissue architecture (13, 14). Investigated at 3, 6 and 12 months (Fig. 2A, top panel), progressive tumor growth in this model is evident macroscopically (Fig. 2A, upper panel), by MR imaging (Fig. 2A, middle panel) and histologically (Fig. 2A, lower panel). Cytokine mRNA expression peaked at 6 months whereas the highest levels of IL1-β, IL-6 and TNF-α were monitored at 12 months (Figs. 2B and C). At earlier time points, the relative expression of IL-6 predominated over the others (Figs. 2C and D), consistent with the potential involvement of IL-6 dependent senescence of mutant cells early on after oncogene activation (22). However, the progressively increased cytokine expression nonetheless indicated that inflammation accompanied tumorigenesis to overt thyroid carcinoma. *Heterogeneous thyroid tumor development features a differential inflammatory response* Since a prominent feature of the sporadic mouse PTC model is multifocal tumor development into distinct PTC subtypes (14, 16), we asked whether cytokine expression might be associated with heterogeneous tumor growth. Indeed, differential although concordant expression levels of *Il1b*, *Il6* and *Tnfa* were evident at 3 months whereas the cytokine expression pattern was highly discordant among individual specimens at 6 months (Fig. 3A). To further characterize oncogene-induced heterogeneity of inflammation, thyroid nodules discerned macroscopically (n=4; Fig. 3B) in a 12 months-old mutant mouse were dissected and individually analyzed for differential gene expression. This showed that *Il1b* and *Il6* transcript levels co-varied in 3 of 4 nodules likely consisting of PTC tumors of different clonal origin (Fig. 3C, upper and middle panels). All tumor foci displayed decreased *Tg* expression varying between 1-19% of that in normal thyroid tissue (Fig. 3C, lower panel), confirming concomitant dedifferentiation of *Braf^CA^*-targeted cells. Histological examination revealed that multifocal thyroid tumors displayed highly variable stromal cell content including lymphocytic infiltration of the immediate tumor microenvironment predominantly found in advanced carcinomas with invasive phenotype (Fig. 4A-D).

**Fig. 2.**
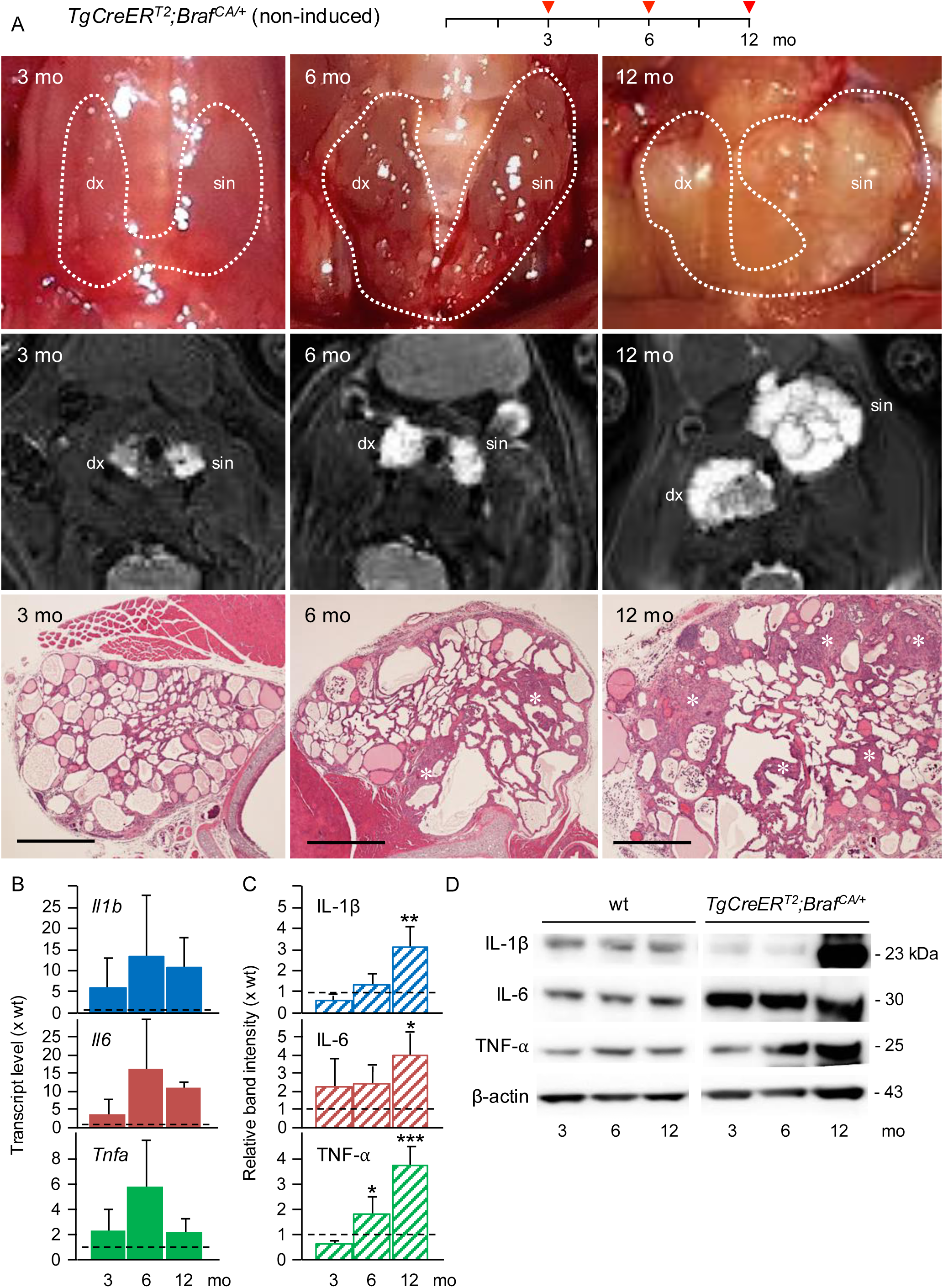
Kinetics of cytokine expression during multifocal tumor development in the thyroid of non-induced *TgCreER^T2^;Braf^CA/+^*mice. Images and data obtained from mutant mice at 3, 6 and 12 months (mo) of age that were devoid of tamoxifen injections. **A** Thyroid gland *in situ* post-surgery (*upper panel*, lobes and connecting isthmus encircled), magnetic resonance imaging (*middle panel*, left (sin) and right (dx) lobes indicated), and histopathology (*lower panel*, midsections of right lobe). Note marked lobe size differences at 12 mo reflecting heterogeneous clonal growth. Asterisks indicate more solid tumor foci. Bars: 500 µm. **B-D** Time-dependent expression changes of IL-1β, IL-6, TNF-α monitored by qPCR (B) and Western blot analysis (C, D). Mean ± SD (qPCT, 3 mo: n=8; 6 mo: n=8; 12 mo: n=6; WB: n=3 for all groups); * p<0.05, ** p<0.01, *** p<0.001. wt wildtype.

**Fig. 3.**
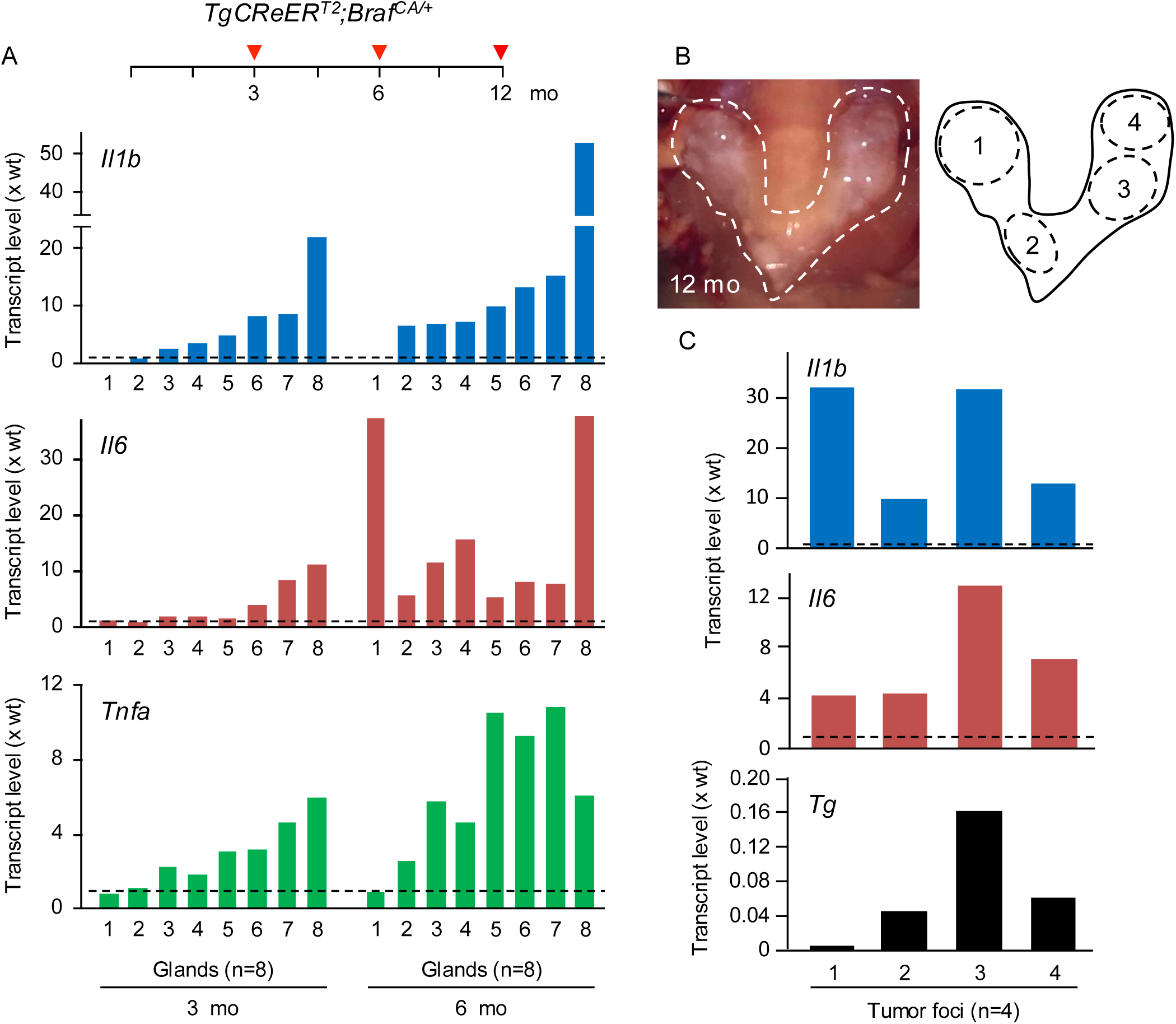
Differential expression of pro-inflammatory cytokines related to multifocal tumorigenesis of BRAF^V600E^-driven papillary thyroid carcinoma. Comparative qPCR data analyzed in tissue samples obtained from non-induced *TgCreER^T2^;Braf^V600E/+^*mice at 3 and 6 months of age (whole lobes) and 12 months old mutants (excised tumors). For all experiment, transcript expression levels were compared to that of control thyroids obtained from age-matched non-mutant (wt) mice. **A** Heterogeneous expression of *Il1b*, *Il6* and *Tnfa* in thyroids of individual mice. Order of presentation deduced by the ranked expression levels of *Il1b* increasing from left to right (n=8) for both time points. . **B-C** Heterogeneous cytokine expression in *Braf^CA^*-induced papillary thyroid carcinoma of different clonal origin. Macroscopically discernable tumors (n=4) were excised from a single mutant thyroid specimen (B, thyroid gland *in situ* (outlined) to the left; cartoon indicating dissected tumor borders to the right) and processed for qPCR analysis of *Il1b* and *Il6*. *Tg* transcripts were simultaneously measured for correlation to level of *Braf^CA^*-induced thyroid dedifferentiation.

**Fig. 4.**
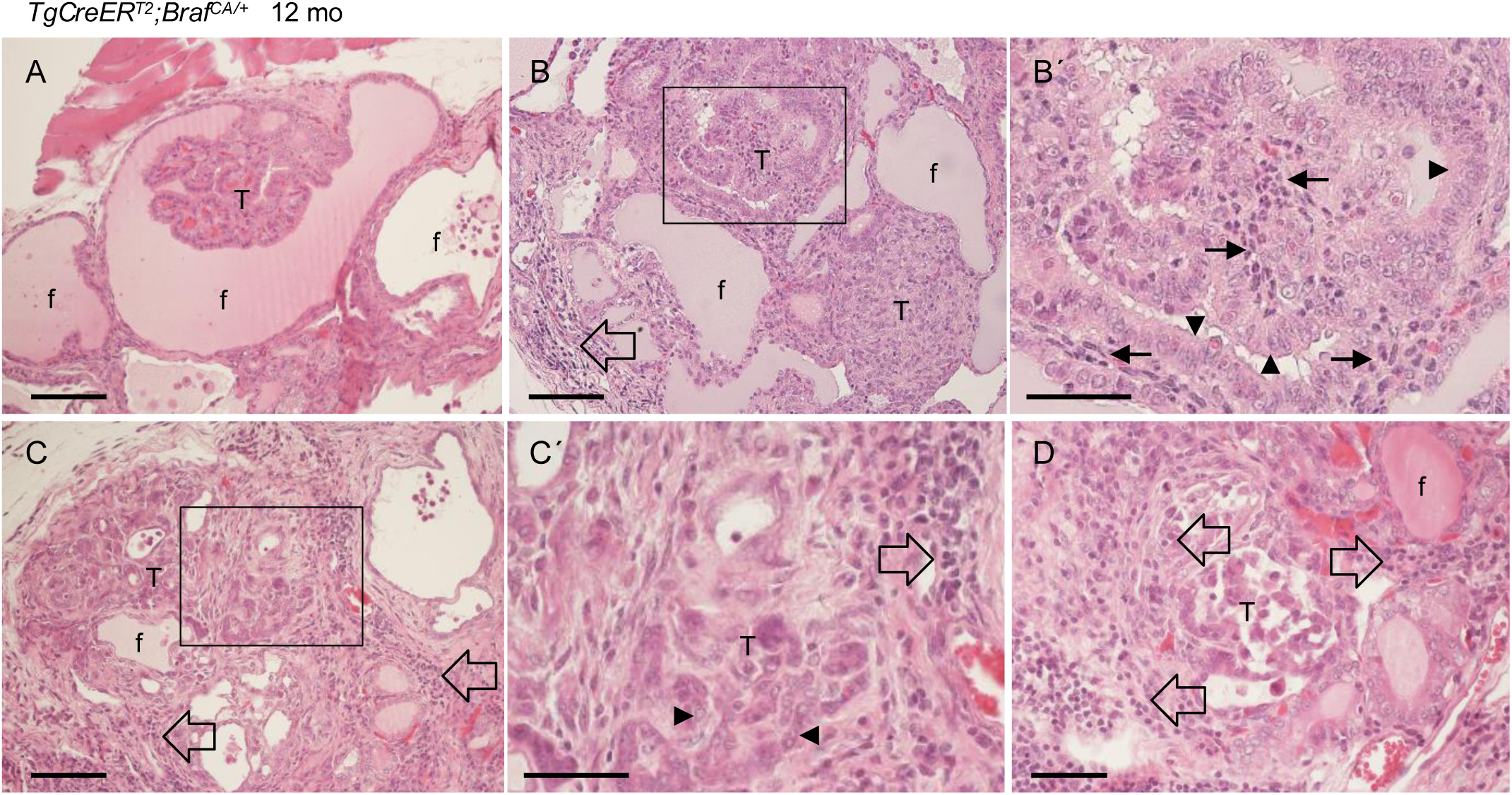
Heterogeneous lymphocytic infiltration of Braf^V600E^-driven thyroid tumors. Representative images of thyroid carcinomas obtained from histological examination of 12 months old non-induced *TgCreER^T2^;Braf^V600E/+^*mice. **A** Papillary thyroid microcarcinoma. **B** Classical and solid variants of papillary thyroid carcinoma. **C, D** Advanced carcinomas with pleomorphic tumor cells. **B′, C′** show high power of boxed area in (B) and (C), respectively. Arrowheads indicate tumor cells, arrows indicate fibrovascular stromal cells, and open arrows indicate clustered lymphocytes. T tumor, f follicle. Scale bars: 100 (A, B, C) and 50 (B′, C′, D) µm.

### Mutant Braf kinase inhibition suppresses Braf^V600E^-induced inflammation along with recovery of thyroid differentiation

Finally, we wanted to investigate reversibility of the inflammatory response to *Braf^CA^* activation. To this purpose, 6 months old non-induced *TgCreER^T2^;Braf^CA/+^* mice were treated daily for 30 days with PLX4720 after which excised thyroids were examined morphologically and for gene expression alterations (Fig. 5, top panel). This showed that drug treatment diminished thyroid lobe size and to large extent restituted thyroid tissue morphology comprising normalized follicle shape and refilling of the follicle lumen with colloid (Fig. 5A, left and right), indicative of redifferentiation of mutant cells. A retained papillary growth pattern of the follicular epithelium identified neoplastic follicles among normal follicles in PLX4720-treated mice (Fig. 5A, right). Transcriptional analysis showed partially recovered expression of thyroid-specific genes implicated in thyroid hormone biosynthesis (*Tg*, *Slc5a5*, *Tpo* and *Tshr*) from <20% to 44-72% of the corresponding expression levels in age-matched wildtype mice (Fig. 5B). *Pax8*, a major transcriptional regulator of thyroid differentiation and function (23–26), showed overshoot expression from ca 50% in untreated to 140% in PLX4720-treated mutant mice (Fig. 5B), presumably reflecting rebound responsiveness of follicular cells with longstanding suppressed thyroid function (27, 28). Notably, PLX4720-treated glands still contained neoplastic follicles with poor colloid content (Fig. 5A, right). Altogether, this suggested that the overall thyroid gene expression levels are the resultant of functionally fully recovered follicles and follicles harboring dedifferentiated non-responding mutant cells.

**Fig. 5.**
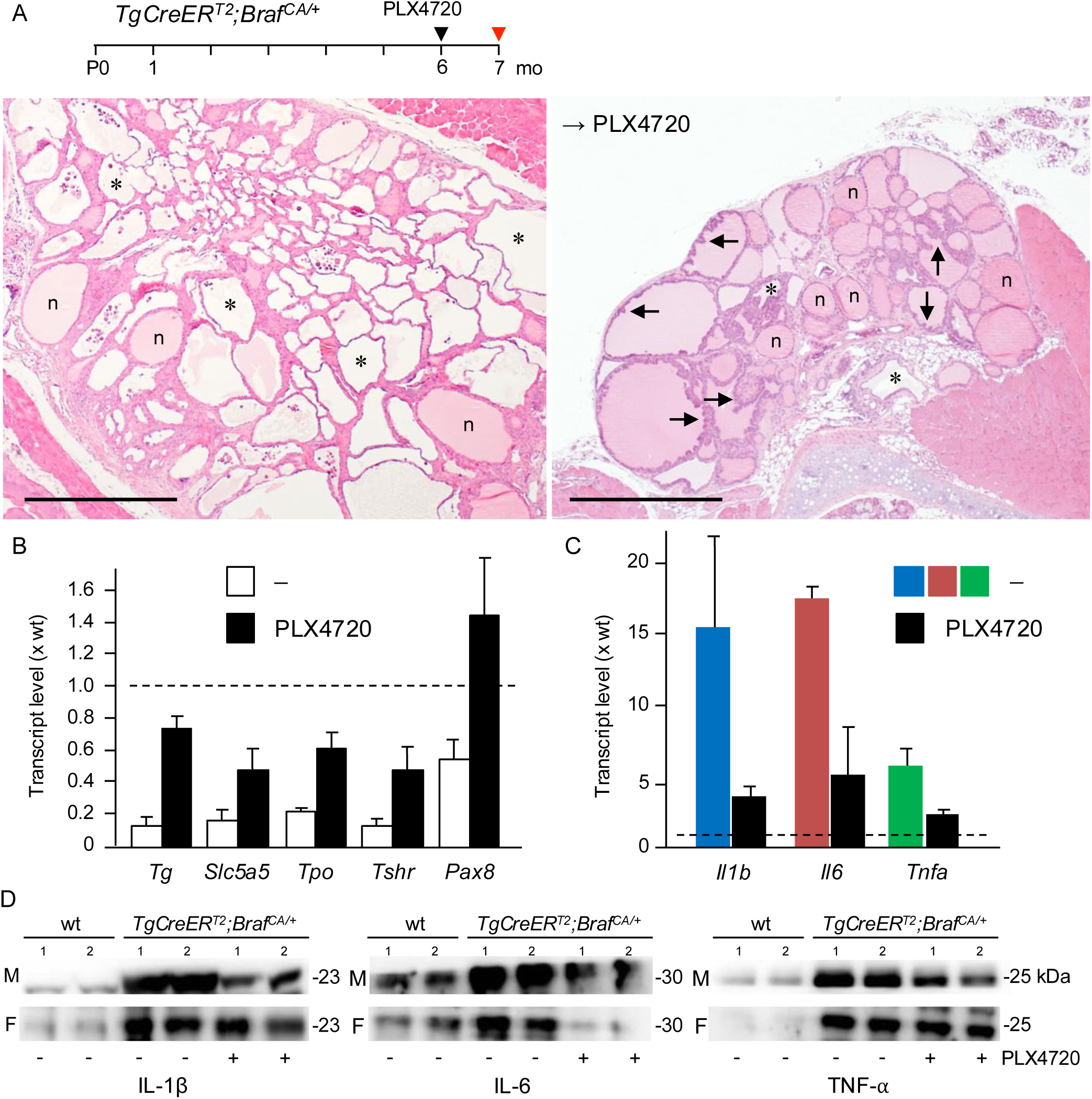
Reduced cancer-associated inflammation accompanies thyroid redifferentiation obtained by mutant BRAF kinase inhibition. Six months old non-induced *TgCreER^T2^;Braf^V600E/+^*mice were treated or not with dietary prodrug to vemurafenib (PLX4720; 417 ppm) for 30 days. Thyroids were processed for morphology, qPCR and Western blot analysis. Transcript expression levels were compared to that of control thyroids in non-mutant mice of the same age. **A** Restitution of follicular morphology by PLX4720. *Left image*: Untreated mutant with thyroid tissue consisting of numerous neoplastic follicles distinguished by loss of colloid designating dedifferentiation. *Right image*: PLX4720-treated mutant with restitution of colloid in most follicles. Note general size reduction of thyroid lobe and many colloid-containing follicles display a papillary growth pattern (arrows). Asterisks indicate follicles with diminished colloid. n normal follicle. Bars: 250 µm. **B** Recovery of thyroid differentiation gene expression in PLX4720-treated mutant mice. *Tg* thyroglobulin, *Slc5a5* sodium iodide symporter (Nis), *Tpo* thyroid peroxidase, *Tshr* thyroid stimulating hormone receptor. Mean ± SD (n=6). **C** Downregulation of cytokine expression in thyroid of PLX4720-treated mutant mice. Samples identical to in B. **D** Heterogeneous anti-inflammatory response to PLX4720. Western blots of thyroid samples from male (M) and female (F) wildtype (wt) and *TgCreER^T2^;Braf^V600E/+^* mice without and with drug treatment (WB: n=3 for all groups); * p<0.05, ** p<0.01, *** p<0.001.

PLX4720 treatment diminished the expression of pro-inflammatory genes (Fig. 5C). However, similarly to the thyroid-specific genes there was a residual expression of *Il1b*, *Il6* and *Tnfa* significantly above the transcript levels in normal thyroids. A heterogeneous response to inhibited BRAFV600E kinase inhibition was also evident in western blot analysis of cytokines in individual mutant glands (Fig. 5D).

## Discussion

This *in vivo* experimental study conceivably recapitulates the time-course of cancer-associated inflammation in BRAF^V600E^-driven PTC. Pro-inflammatory cytokine expression comprising Il-1ß, Il-6 and Tnf-α is evident in the mutant follicular epithelium already before tumorigenesis is initiated. The level of cytokine-based inflammation increases as tumors progress to microcarcinomas and overt PTC. The cytokine expression pattern is rather uniform in early tumor development whereas the cytokine signature is highly variable among individual tumor foci in advanced tumor stages. Cytokine expression levels diminish by inhibition of mutant Braf kinase indicating that inflammation is a direct consequence of constitutive activation of the MAPK signaling pathway.

BRAFV600E is the most prevalent driver mutation in thyroid cancer primarily encountered in PTC, although the same BRAF mutant allele is also frequently encountered in anaplastic thyroid carcinoma (ATC) presumably designating tumor progression from PTC to ATC (29). Indeed, recent tumor DNA sequencing indicate that ATC share a genomic and evolutionary origin with co-occurring differentiated thyroid carcinomas featuring additional acquired driver mutations (30). There are currently no animal models that recapitulate PTC-ATC transition involving sequentially increased mutational burden. Co-expressed BRAF^V600E^ and PIK3CA^H1047R^ or homozygous inactivation conditionally of *TP53* or *PTEN* accompanying *Braf^CA^* activation confer tumor progression to an ATC-like phenotype (31, 32), however with a latency of six months or more suggesting that yet unidentified factors presumably accumulating over time related to tumor growth contribute to ATC pathogenesis. Characterization of the tumor immune microenvironment has revealed that immune evasion is a prominent feature of ATC (5). Since constitutive PI3′kinase signaling or loss of p53 or Pten expression are not per se tumorigenic in mouse ATC models (31, 32), it is presumed that BRAF^V600E^ drives immunosuppression eventually beneficial to tumor progression.

Our present findings indicate that mutant Braf elicits both acute and chronic inflammation, the former likely associated with oncogene-induced senescence whereas the delayed response accompanies tumor development to microcarcinomas of which some transition to overt PTC. Differential expression of pro-inflammatory cytokines is attributed to distinct tumor foci and variably among age-matched mutant mice, strongly suggesting that oncogene-induced inflammation is a clonal trait. We previously showed that spontaneous *Braf^CA^*activation due to leaky Cre activity in the current model reproduces the typical heterogeneous tumor phenotypes that characterize PTC subtypes and that clonality of tumor development can be traced back to the originating thyroid follicle (14). Normal follicles are not uniform but differ much in terms of both cell content and lumen compartment, and individual cells differ in gene expression (33) and responsiveness to mitogenic stimuli (34). We hypothesize that the natural follicle heterogeneity, which primarily relies on clonal segregation of progenitor cells during embryonic thyroid development (35), constitute the basis of tumor heterogeneity (16), in a sense homologous to the pathogenesis of multinodular goiter (36, 37). Thus, the fate of a Braf mutant thyroid cell is at least partly determined by intrinsic properties of the affected follicle, for example, whether outcompeted by non-mutant cells or synergizing with another mutant clone (14). A heterogeneous inflammatory response to oncogenic activation as reported here might have a similar histoarchitectural foundation.

Consistent with a documented role of IL-6 in oncogene-induced senescence (22), enhanced thyroid expression of Il-6 predominated initially in both induced and non-induced mutant mice. However, sustained expression of Il-6 might contribute to thyroid tumorigenesis and progression as suggested from previous studies on human PTC tumor cells (38, 39). Likewise, both IL-6 and IL-1ß expression are increased in Braf mutant melanoma cells (40). In FRTL-5 cells, an immortalized differentiated rat thyroid cell line, IL1ß stimulates IL-6 expression (41). Although these observations most likely are cell type and context-dependent, it is conceivable to assume that cytokines secreted in response to stochastic *Braf^CA^*activation primarily derive from mutant thyroid cells, as supported by our RNAscope findings. By contrast, induced *Braf^CA^* activation led to rapid accumulation of *Il1ß*^+^ cells and to a lesser extent *Il6*^+^/*Tnfa*^+^ cells to the stromal compartment, thus deviating from the expression pattern associated with sporadic tumorigenesis. Notably, both models confer oncogene-induced senescence, but whether senescence arises within incipient tumor cells or stromal cells might differentially impact on tumorigenesis through the senescence-associated secretory phenotype (SASP) (42). An additional level of complexity is inferred by recent findings of an intracrine mechanism of IL-6-mediated senescence (43).

Accompanying restitution of follicular histoarchitecture and recovery of thyroid gene expression, PLX4720 inhibited Braf^V600E^-mediated expression of all three cytokines. It is previously known that both IL-1α and IL-6 suppresses thyroid function in cultured normal thyroid follicular cells (44, 45). IL-1 (both alpha and beta) but not IL-6 negatively influences the thyroid epithelial barrier by modifying tight junctions inferring a role of paracellular leakage in thyroid autoimmunity (46–48). Moreover, in vitro-reconstituted thyroid follicles show lumen dilation and flattened epithelium in response to IL-1α (45), which is reminiscent of the general morphological changes that feature many follicles in untreated *TgCreER^T2^;Braf^CA/+^* mice (this study and (14)). It is thus possible that recovery of thyroid function and follicular structure in response to mutant Braf kinase inhibition might at least partly be mediated by diminished cytokine action.

Since PLX4720 only partially restituted thyroid gene expression (except for overshooting Pax8), it is possible that the residual overexpression of cytokines might reflect drug resistance. Importantly, drug treatment was evaluated in Braf mutant mice between 6 and 7 months, which correspond to an intermediate microcarcinoma stage of PTC development. Moreover, transcript quantification was performed on whole lobe tissues consisting of a mixture of normal follicular cells and mutant cells of which only some clones are tumorigenic (14). Therefore, it is likely that maintained cytokine expression in drug-treated mice is attributed to a minor fraction of Braf mutant thyroid cells or, alternatively, the stromal microenvironment. Both possibilities gain support from previous studies in melanoma cells as spliced variants of IL-6 confer resistance to vemurafenib by reactivating the MAPK pathway (49) whereas Braf kinase inhibitors may also directly stimulates IL-1ß secretion by dendritic cells presumed to be part of the anti-tumor immune response (50). Presence of absence of drug-resistant clones – primarily related to stochastic *Braf^CA^* activation multifocally – probably explain the variable efficiency of Braf kinas inhibition on cytokine expression among thyroid glands from both sexes. A putative clone-bias of differential drug responsiveness adds another level of tumor heterogeneity in Braf^V600E^-driven thyroid cancer.

## Conclusions

In a mouse model of sporadic PTC development, BRAF^V600E^ driver mutation confers a heterogeneous pro-inflammatory response indicating that the tumor immune microenvironment is clonally determined. In view of previous findings that concurrent secondary mutations are rare even in advanced tumor stages (14), these findings are consistent with the hypothesis that different tumor-cell-origin properties inherent to the natural follicular heterogeneity imprint on tumor development and ultimately the tumor phenotype comprising also immunogenicity. Tumorous cytokine production is a direct consequence of mutant Braf kinase activity. Drug resistance to Braf kinase inhibition as revealed by sustained inflammation is recognized already in some subclinical tumors corresponding to papillary microcarcinomas.

## List of abbreviations

PTC: papillary thyroid carcinoma
MAPK: mitogen-activated protein kinase
Tg: thyroglobulin
Tam: tamoxifen
Tpo: thyroid peroxidase
Tshr: thyroid stimulating hormone receptor
IL: interleukin
TNF: tumor necrosis factor
ISH: in situ hybridization
FFPE: formalin-fixed paraffin-embedded
WB: Western blot
HRP: horseradish peroxidase
IHC: immunohistochemistry
MR: magnetic resonance
qRT-PCR: quantitative real-time polymerase chain reaction
ATC: anaplastic thyroid carcinoma
SASP: senescence-associated secretory phenotype
Nis: sodium iodide symporter

## Declarations

### Ethic statement

This animal study was approved by Göteborgs Djurförsöksetiska Nämnd (5.8.18-04502/2023) according to European standards and national regulations provided by the Swedish Board of Agriculture.

### Concent for publication

not applicable.

### Availability of data and materials

The datasets used and analyzed during the current study are available from the corresponding author (MN) on reasonable request.

### Authors′ contributions

MN conceptualized the study and wrote the manuscript draft; SK performed Western blot and RNAScope experiments; CM was responsible for qRT-PCR analysis; HF contributed to the interpretation of data as clinical pathologist; ES was responsible for all animal experiments; all authors read and approved the final manuscript.

## Acknowledgements

not applicable.

