## Supplemental data for "Heterogeneous pro-inflammatory response to BRAFV600E-induced thyroid tumor development"

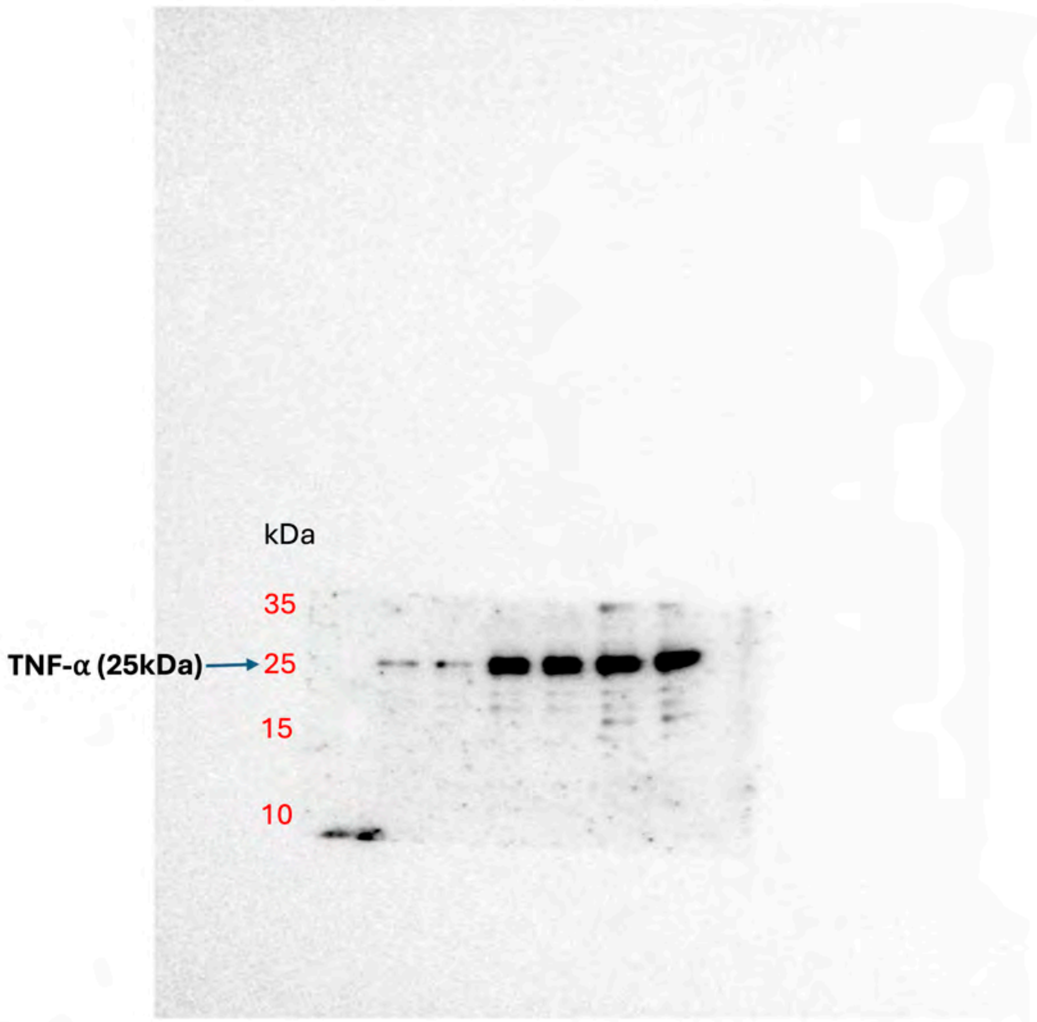


Figure 1 (C) : Uncropped blot for TNF a. Molecular weight markers indicated. **Ladder:** PageRuler™ Plus Prestained Protein Ladder (Thermo Fisher, Cat# 26619). Membrane was cut prior to antibody incubation and developed separately.

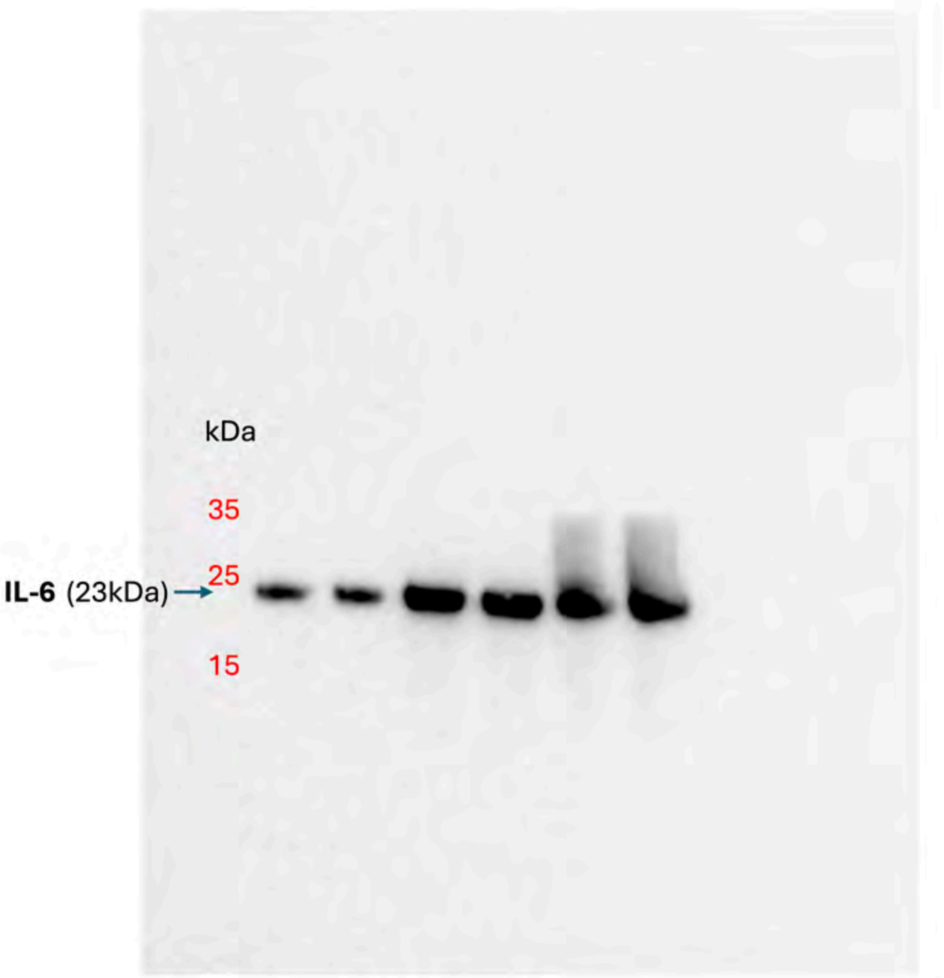


Figure 1 (C) : Uncropped blot for IL6. Molecular weight markers indicated. **Ladder:** PageRuler™ Plus Prestained Protein Ladder (Thermo Fisher, Cat# 26619). Membrane was cut prior to antibody incubation and developed separately.

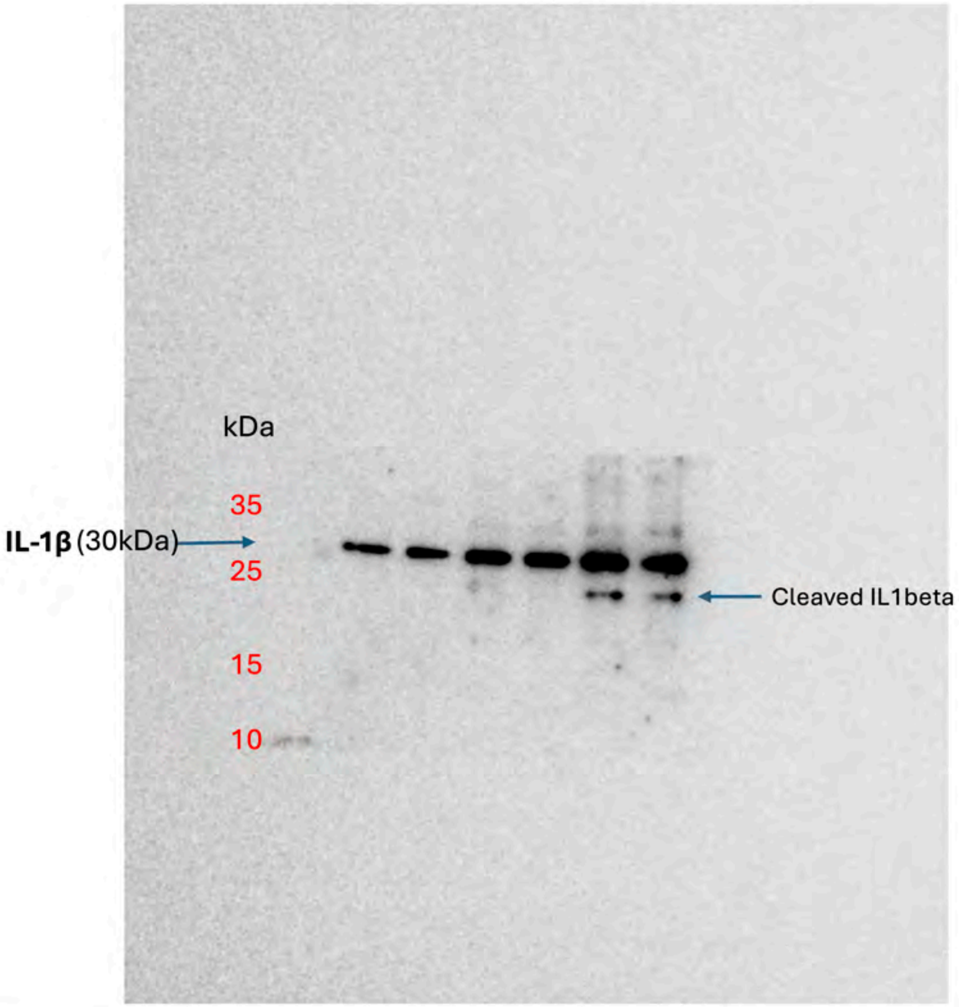


Figure 1 (C) : Uncropped blot for IL-1β. Molecular weight markers indicated. **Ladder:** PageRuler™ Plus Prestained Protein Ladder (Thermo Fisher, Cat# 26619). Membrane was cut prior to antibody incubation and developed separately.

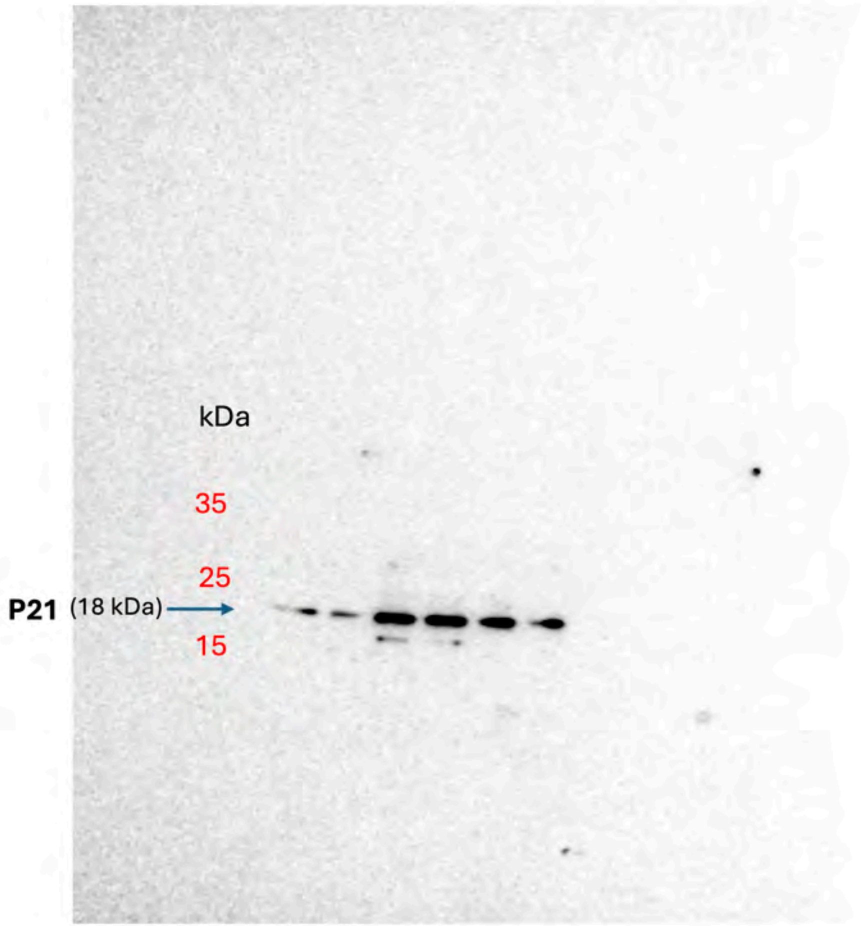


Figure 1 (C) : Uncropped blot for P21. Molecular weight markers indicated. **Ladder:** PageRuler™ Plus Prestained Protein Ladder (Thermo Fisher, Cat# 26619). Membrane was cut prior to antibody incubation and developed separately.

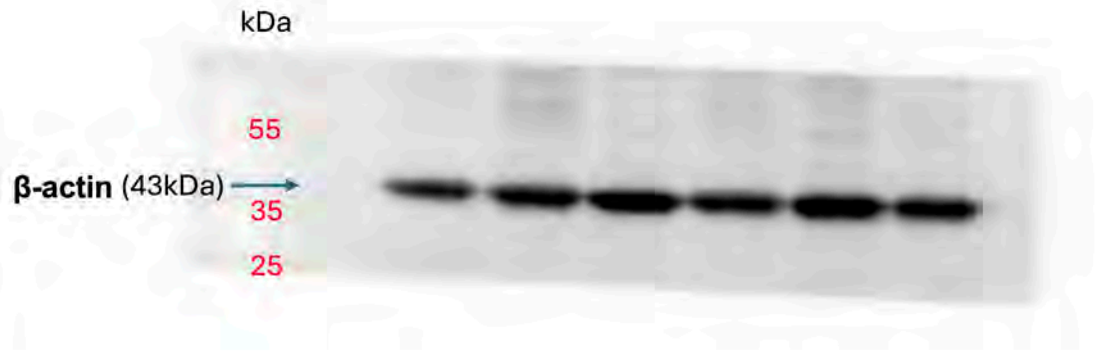


Figure 1 (C) : Uncropped blot for β-actin. Molecular weight markers indicated. **Ladder:** PageRuler™ Plus Prestained Protein Ladder (Thermo Fisher, Cat# 26619). Membrane was cut prior to antibody incubation and developed separately.

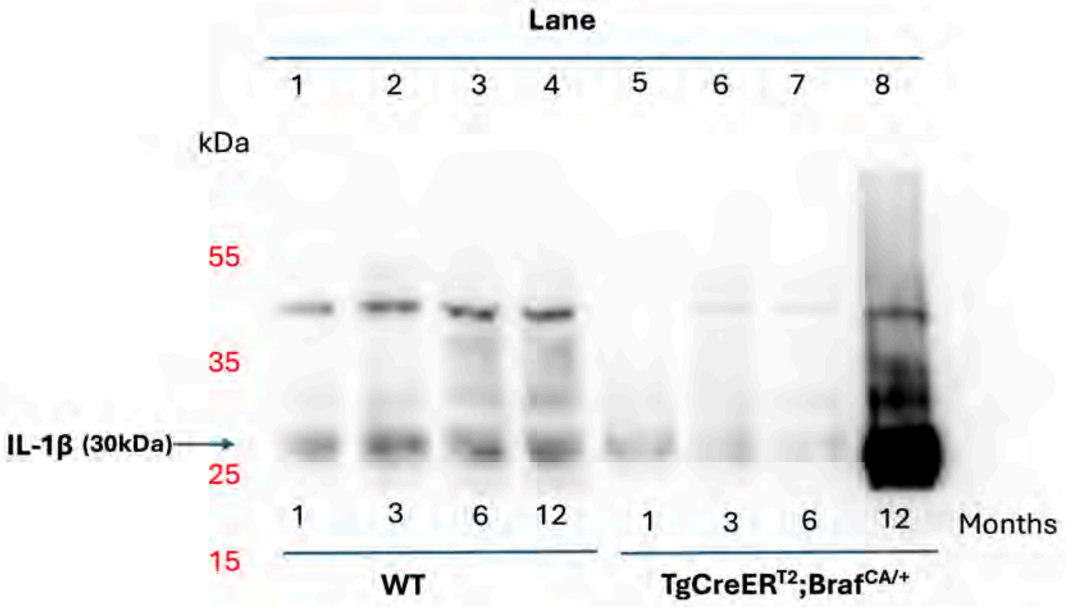


**Figure 2 (D)**: Uncropped blot for IL-1β. Molecular weight markers indicated. **Ladder:** PageRuler™ Plus Prestained Protein Ladder (Thermo Fisher, Cat# 26619).
Lanes corresponding to 1-month samples (lanes 1 and 5) were omitted from the main Figure 2 to align with the MRI-based analysis focusing on the 3-, 6-, and 12-month time points. Membrane was cut prior to antibody incubation and developed separately.

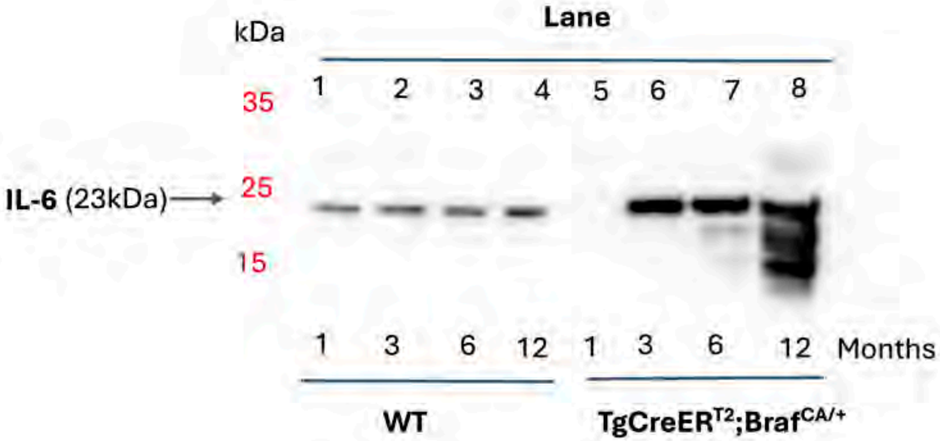


**Figure 2 (D)**: Uncropped blot for IL-6. Molecular weight markers indicated. **Ladder:** PageRuler™ Plus Prestained Protein Ladder (Thermo Fisher, Cat# 26619). Lanes corresponding to 1-month samples (lanes 1 and 5) were omitted from the main Figure 2 to align with the MRI-based analysis focusing on the 3-, 6-, and 12-month time points. Membrane was cut prior to antibody incubation and developed separately.

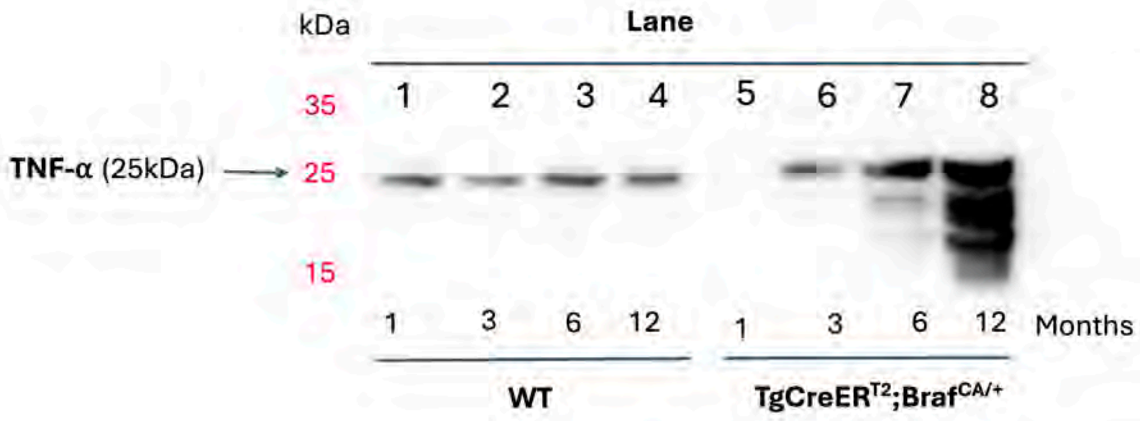


**Figure 2 (D)**: Uncropped blot for TNF a. Molecular weight markers indicated. **Ladder:** PageRuler™ Plus Prestained Protein Ladder (Thermo Fisher, Cat# 26619). Lanes corresponding to 1-month samples (lanes 1 and 5) were omitted from the main Figure 2 to align with the MRI-based analysis focusing on the 3-, 6-, and 12-month time points. Membrane was cut prior to antibody incubation and developed separately.

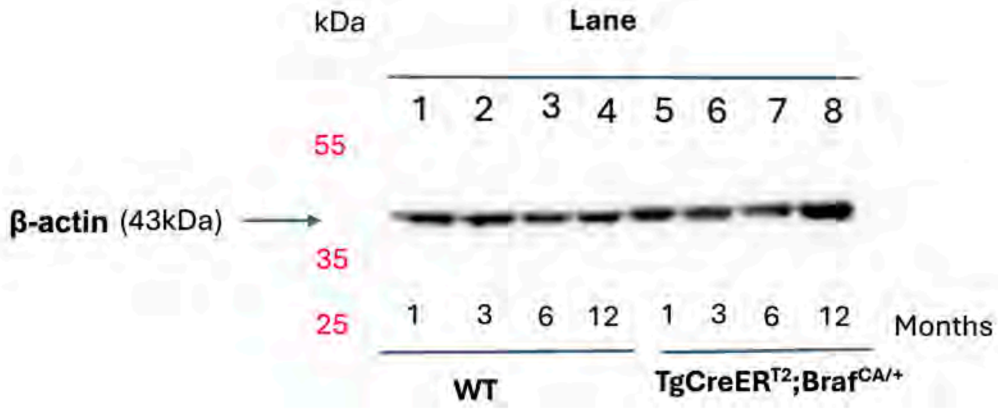


**Figure 2 (D)**: Uncropped blot for β-actin. Molecular weight markers indicated. **Ladder:** PageRuler™ Plus Prestained Protein Ladder (Thermo Fisher, Cat# 26619). Lanes corresponding to 1-month samples (lanes 1 and 5) were omitted from the main Figure 2 to align with the MRI-based analysis focusing on the 3-, 6-, and 12-month time points. Membrane was cut prior to antibody incubation and developed separately.

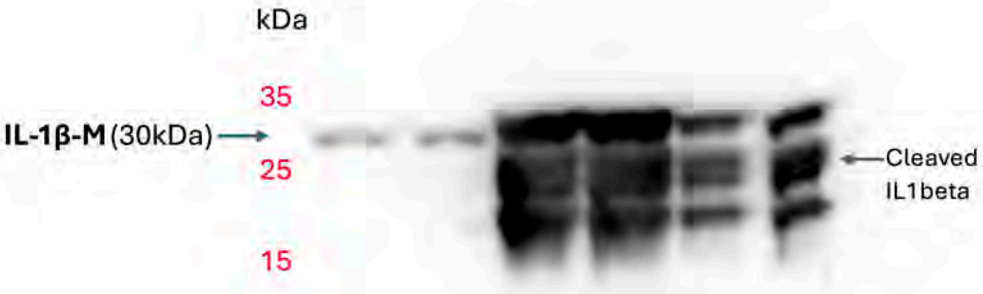


**Figure 5 (D)**: Uncropped blot for ILbeta-M. Molecular weight markers indicated. **Ladder:** PageRuler™ Plus Prestained Protein Ladder (Thermo Fisher, Cat# 26619). Membrane was cut prior to antibody incubation and developed separately.

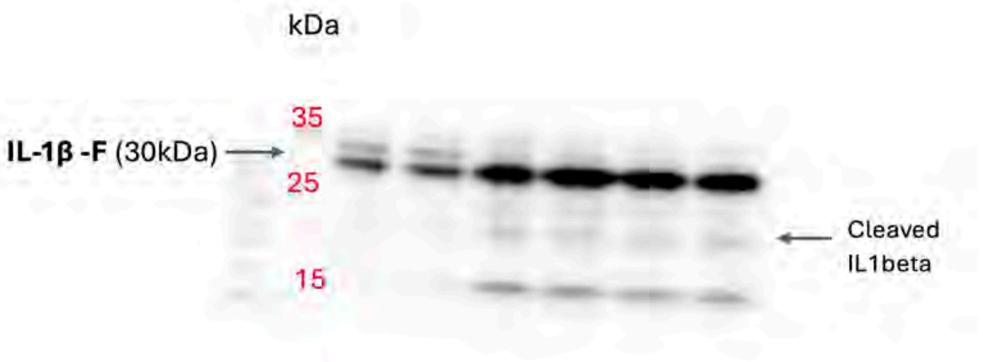


**Figure 5 (D)**: Uncropped blot for ILbeta-F. Molecular weight markers indicated. **Ladder:** PageRuler™ Plus Prestained Protein Ladder (Thermo Fisher, Cat# 26619). Membrane was cut prior to antibody incubation and developed separately.

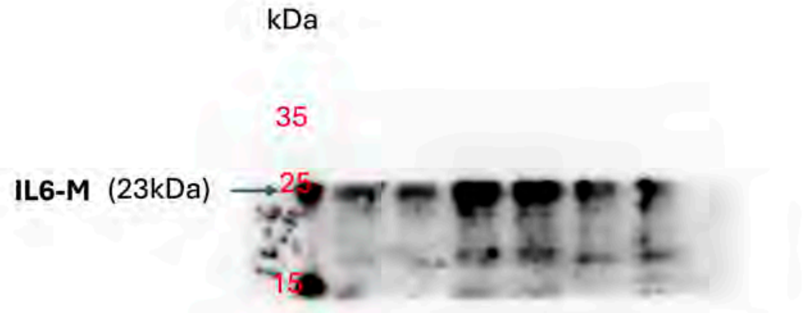


**Figure 5 (D)**: Uncropped blot for IL-6-M. Molecular weight markers indicated. **Ladder:** PageRuler™ Plus Prestained Protein Ladder (Thermo Fisher, Cat# 26619). Membrane was cut prior to antibody incubation and developed separately.

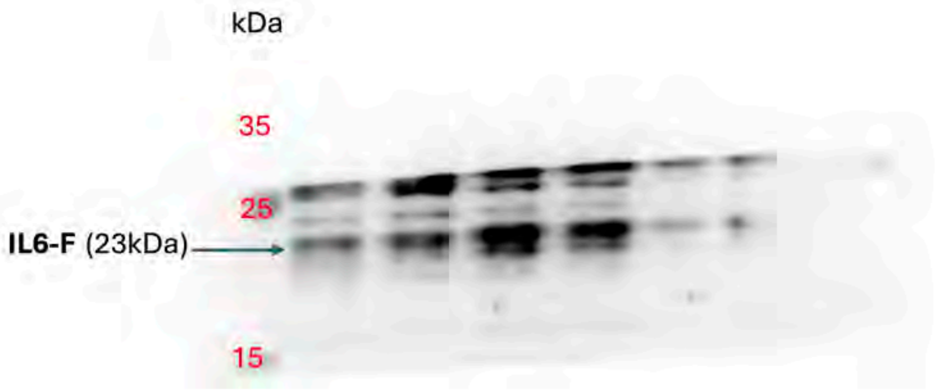


**Figure 5 (D)**: Uncropped blot for IL-6-F. Molecular weight markers indicated. **Ladder:** PageRuler™ Plus Prestained Protein Ladder (Thermo Fisher, Cat# 26619). Membrane was cut prior to antibody incubation and developed separately.

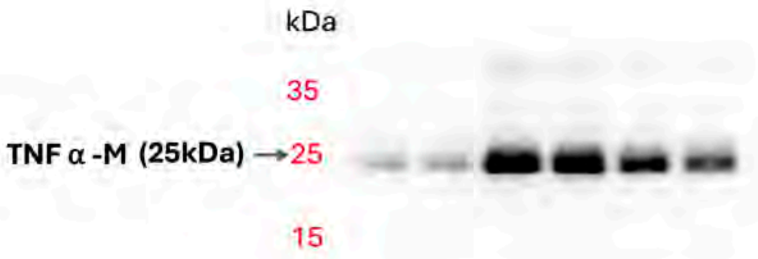


**Figure 5 (D)**: Uncropped blot for TNF a-M. Molecular weight markers indicated. **Ladder:** PageRuler™ Plus Prestained Protein Ladder (Thermo Fisher, Cat# 26619). Membrane was cut prior to antibody incubation and developed separately.

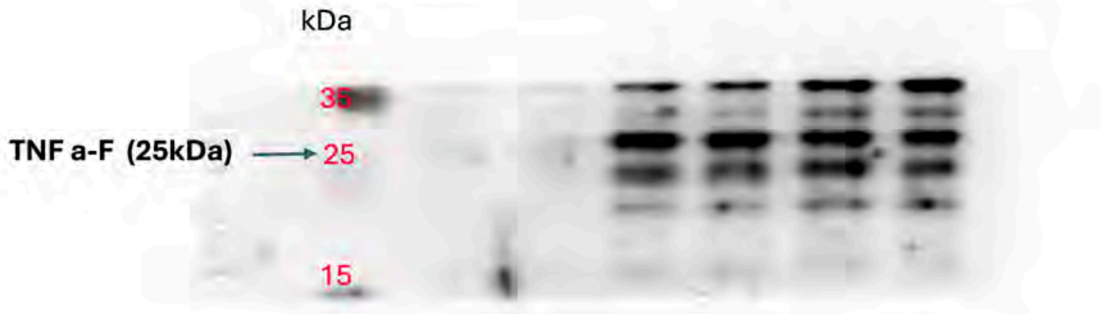


**Figure 5 (D)**: Uncropped blot for TNF a-F. Molecular weight markers indicated. **Ladder:** PageRuler™ Plus Prestained Protein Ladder (Thermo Fisher, Cat# 26619). Membrane was cut prior to antibody incubation and developed separately.
